# Structural connectivity analysis using Finsler geometry

**DOI:** 10.1101/424150

**Authors:** Tom Dela Haije, Peter Savadjiev, Andrea Fuster, Robert T. Schultz, Ragini Verma, Luc Florack, Carl-Fredrik Westin

## Abstract

In this work we demonstrate how Finsler geometry—and specifically the related geodesic tractography— can be levied to analyze structural connections between different brain regions. We present new theoretical developments which support the definition of a novel Finsler metric and associated connectivity measures, based on closely related works on the Riemannian framework for diffusion MRI. Using data from the Human Connectome Project, as well as population data from an autism spectrum disorder study, we demonstrate that this new Finsler metric, together with the new connectivity measures, results in connectivity maps that are much closer to known tract anatomy compared to previous geodesic connectivity methods. Our implementation can be used to compute geodesic distance and connectivity maps for segmented areas, and is publicly available.

## 1. Introduction

Finsler geometry was proposed as a means to analyze diffusion MRI data in works by Pichon et al. [36] and Melonakos et al. [31, 32], where the authors computed shortest geodesic tracts based on an ad hoc relation with the diffusion MRI signal. A different definition of the Finsler function was employed by Astola et al. [4], who illustrate some more technical applications of Finsler geometry, while the work of Sepasian et al. [41] extends shortest geodesic tractography to allow multiple geodesic connections between points. Finally, the first geodesic connectivity analyses were performed by de Boer et al. [13], which also marks the first time Finsler geometry was used to study group differences in diffusion MRI data. Though these results demonstrate the potential value of Finsler geometry in diffusion MRI, one downside is the limited availability of tools that can perform these types of analyses. For this reason, along with the highly technical content of even introductory works on Finsler geometry, Finsler-based methods remain inaccessible to the majority of diffusion MRI researchers.

The goal of this paper is to provide an informal introduction to the core concepts involved with Finsler connectivity computations, based on our recently released Finsler connectivity module (FCM). This module is an adaptation of the `finslertract` code by Antonio Tristán-Vega (nitrc.org/projects/finslertract), which is used to perform Finsler tractography. To this end, we describe the ideas behind the application of Finsler geometry to diffusion-weighted MRI data in section 2, and describe the tractography and connectivity algorithm in section 3. This section introduces a new Finsler metric definition and a generalization of a connectivity measure used for diffusion tensor imaging [6], in order to address theoretical limitations of existing geometrical methods. As a practical illustration of the method, we compute the connectivity for a number of well-known major fiber bundles in a high resolution public data set. We also discuss how the connectivity algorithm can be used in group studies and, as a proof of concept, use our method to study group differences in autism spectrum disorder data using network-based analysis techniques. The results of these experiments are presented in section 4. Finally we discuss some strengths and shortcomings of the approach in section 5.

## 2. Theoretical background

### 2.1. Finsler geometry

Finsler geometry is concerned with measuring distances on abstract spaces called Finsler manifolds. A Finsler manifold consists of a base space, in the context of this paper always the set of positions in ℝ^3^ where diffusion is measured, and a real scalar-valued function *F*: ℝ^3^ × ℝ^3^ → ℝ^+^ that captures the additional structure of the space. The distance between two points on a Finsler manifold is defined similar to the standard Euclidean distance, namely in terms of the length of the shortest curve connecting the two points; the difference lies in the way the length of the curve is computed. The length 𝓛_*F*_(*γ*) of a curve *γ*: [0, *L*] → ℝ^3^ is still a sum over infinitesimal line elements d*γ*, but the associated length of each line element is now weighted depending on both its position and its orientation:

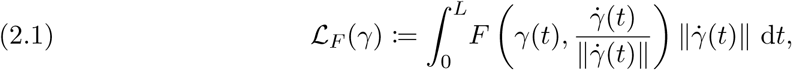

where 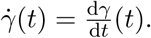 With *F* = 1 this reduces to the Euclidean length of the curve,

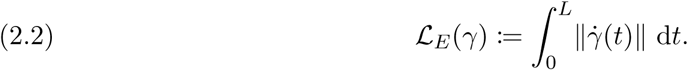

The Finsler function *F* is (positively) homogeneous of degree one in its second argument, which means that for any point **x** and any direction **y** the following relation holds:

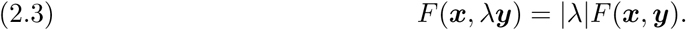

We can thus write

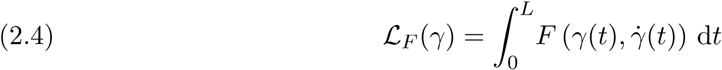

and

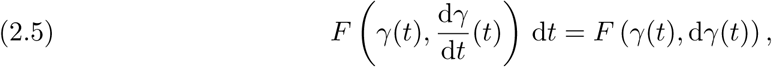

which clarifies the interpretation of *F* as a function acting locally on a line element d*γ*. At the same time we note that the homogeneity of *F*, (2.3), guarantees that the length *F* (*γ* (*t*), d*γ* (*t*)) associated to d*γ* is determined solely by its orientation—not by its ‘magnitude’—and that 𝓛_*F*_ (*γ*) is independent of the (proper) parametrization of *γ*. As a technical aside we observe that homogeneity also implies that *F* is strictly speaking defined only when 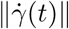≠ 0 for all *t*, and for this reason we have to assume a parametrization of *γ* that avoids this issue. To simplify the following discussion we assume without loss of generality that 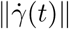= 1, i.e., we assume that 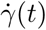 is an element of the sphere S^2^. Some additional properties and technicalities related to *F* are given in the paper by Florack et al. [18], and a thorough exposition on Finsler geometry can be found for example in the book by Bao et al. [5].

### 2.2. The Finslerian framework for diffusion MRI

The Finslerian framework for diffusion MRI models the brain as a Finsler manifold, by deriving a Finsler function *F* from diffusion-weighted data. The principal idea behind this is the well-known correlation between the local amount of diffusion in a certain direction, and the large scale structural orientation of white matter [8]. By defining (or assuming) a correspondence between the Finslerian length of a curve and the amount of diffusion along a curve, we can leverage a rich set of Finsler geometrical tools for the analysis of diffusion data.

Bearing this in mind, the Finsler function is generally defined such that some measure of diffusivity (e.g. a diffusion orientation distribution function [44], or dODF) at a given point and in a certain direction, is inversely related to the associated length. In other words, we have that a large diffusivity at a point ***x*** along a vector ***y***, corresponds to a small value *F*(***x***, ***y***). This leads to the useful alternative viewpoint of the Finsler function as a kind of cost function. If we consider a displacement in direction ***y*** as a parameter that can be controlled, then *F* can be interpreted as associating a high cost to movement in a direction with low diffusivity, and a low cost to movement in directions of high diffusivity.

### 2.3. Geodesics

A prime example of analysis tools made available by the geometrical framework are geodesics. Geodesics can be regarded as connections along which one en-counters, in some sense, optimal diffusivity. More specifically, geodesics connecting two given points are those curves for which the length 𝓛_*F*_ is (locally) minimal, and can thus be viewed as the Finslerian analogue of ‘straight lines’. The existence of a geodesic between any two points in the Finsler manifold is guaranteed [5, Chapter 6.6], which means that we can find optimal connections between any two points or regions of interest. For now we will assume that the shortest geodesic between two points, called the minimal geodesic, is uniquely defined.

In practice we determine minimal geodesics using a fast-sweeping algorithm [31] based on the principle of optimality, which states that given a unique minimal geodesic *γ*: [0, *L*] → ℝ^3^, the geodesic segment between the points *γ*(*a*) and *γ*(*b*), *a*,*b* ∈ [0, *L*], is necessarily identical to the geodesic between these two points. If we write 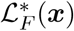 for the shortest geodesic distance from a point ***x*** ∈ ℝ^3^ to a seed region Ω ⊂ ℝ^3^ relative to *F*, then

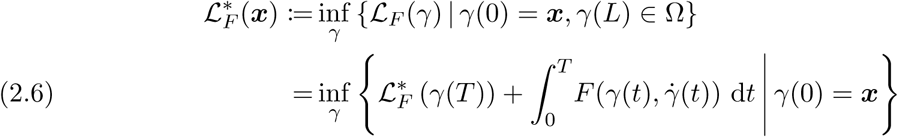

for all *T* ∈ (0, *L*), cf. Figure 1. From the Taylor expansion

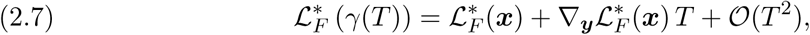

where ∇_***y***_ denotes the directional derivative along ***y*** and where we use the shorthand notation ***x*** = *γ*(0) and ***y*** =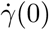 ∈ S^2^, we find the Hamilton–Jacobi–Belmann equation in the limit *T* → 0:

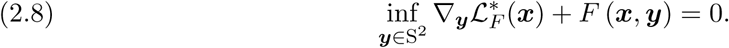

**Figure 1.**
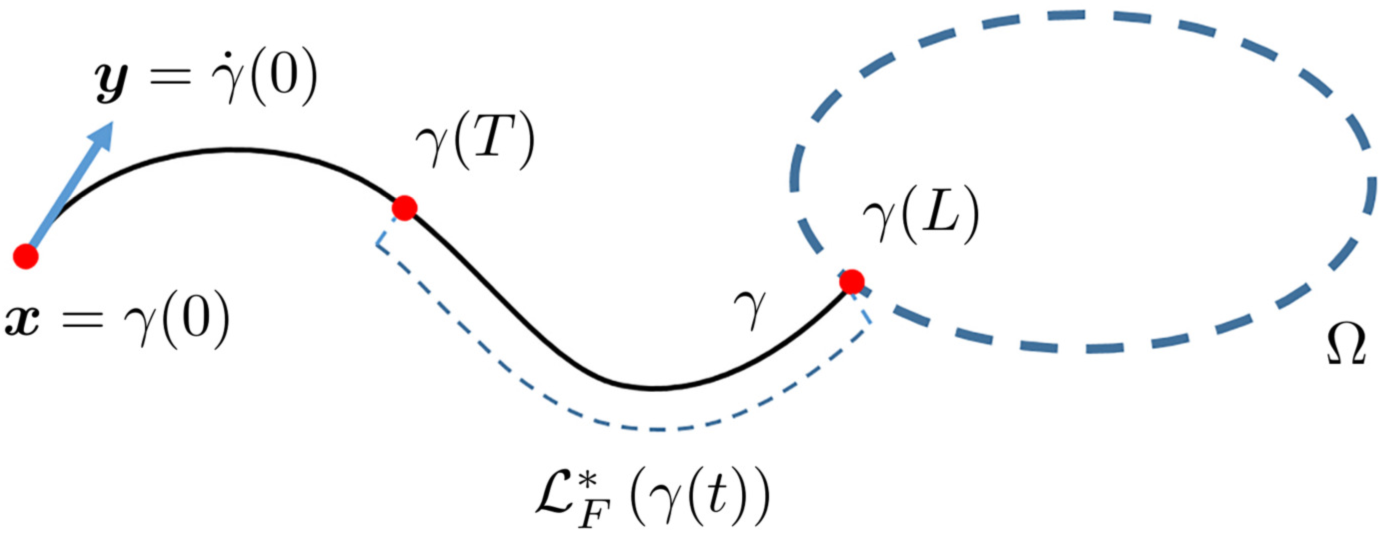
The principle of optimality states that segments of the minimal geodesic between two points are themselves geodesic. The black curve represents the optimal curve γ (minimal geodesic) connecting the point x to the seed region Ω, i.e., the curve γ that minimizes the Finslerian length functional 𝓛_F_. The distance 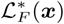 from x to Ω is defined as the length of the optimal curve γ that connects the two. If 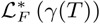 is known for γ(T) near x, the principle of optimality allows us to compute 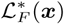 by solving the Hamilton–Jacobi–Bellman equation (2.8). As 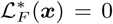 for all x ∈ Ω, repeated application of (2.8) allows us to compute 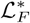 for all x ∈ M.

Together with the initial condition

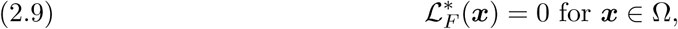

repeated application of (2.8) allows us to compute the complete 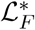 map for all ***x*** ∈ ℝ^3^. See Appendix A for implementation details.

This method is relatively fast, but as explained before it is limited in that it only finds the shortest geodesic out of the possibly many geodesics connecting two given points [41]. In the following we will make the usual assumption that all relevant information is captured by the minimal geodesic. More information on fast-sweeping algorithms can be found in the references [25, 24, 48].

Returning to the context of diffusion MRI, the optimal diffusivity along geodesics is made more precise by the inverse relation discussed in subsection 2.1. With the cost function interpretation of *F*, we note that geodesics correspond to curves along which the accrued cost is minimal. Because of the inverse relation between the cost and the local diffusivity, geodesics thus minimize movement in directions of low diffusivity. Additionally, we see that the Finslerian length 𝓛_*F*_ of a curve approximately corresponds to the average reciprocal diffusivity along the curve.

## 3. Methods

Geodesics have been used as a tool in tractography, based on the hypothesis that (some small subset of) geodesics between two points coincide with the physical connections between them. This approach has certain practically advantageous features; it relies on the full diffusion information available (unlike most deterministic streamline tractography algorithms for instance), and has essentially no parameters that have to be tuned. It should however be noted that since there is no canonical definition of the Finsler function, this choice of ‘metric’ does add an additional degree of freedom to the algorithm. The `riemanntract` and `finslertract` packages available on nitrc.org can be used to perform geodesic tractography with the different metrics.

Geodesic connectivity analysis is based on the hypothesis that curves of optimal diffusivity between two points reflect the likelihood of a structural connection. Although this means that high connectivity values are expected within anatomical bundles, this does not imply that geodesic curves necessarily trace existing physical connections as they should in the case of geodesic tractography. In this section we discuss the options for the Finsler function in the FCM, as well as two available geodesic-based path and connectivity measures that have been studied in recent works.

### 3.1. The Finsler function

The choice for the Finsler function *F* in terms of the diffusion signal is the primary degree of freedom in the Finslerian framework, and determines to a large extent how geometric features such as geodesics can be interpreted. The most common choice for *F* found in the literature was proposed by Melonakos et al. [32] and is given by

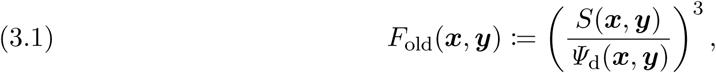

with ***y*** ∈ S^2^ and *S* the diffusion MRI signal acquired on a fixed *b*-value shell, and where *Ψ*_d_ is the diffusion orientation distribution function (dODF) defined in terms of the Funk–Radon transform (see e.g. Tuch et al. [44]). The power 3 is used as a type of sharpening. The choice *F* = *F*_old_ has been shown to produce reasonable tractography results e.g. near the cingulum bundle [31], and it has been used in the literature by de Boer et al. [13]. However, it lacks a clear relation with the more well-founded existing metrics in the Riemannian framework—we expect the Finsler function in case of Gaussian diffusion to reduce to a Riemannian norm compatible with the canonical Riemannian metric [34, 27], or one of the metrics derived therefrom [22, 20, 23, 19].

As an alternative we postulate a new choice for *F*:

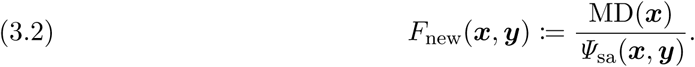

where MD is a generalization of the mean diffusivity used in diffusion tensor imaging (DTI) [7] defined as the average apparent diffusion coefficient [47], and *Ψ*_sa_ is the solid angle dODF [1]. In Appendix B we show that for purely Gaussian diffusion, *F*_new_ corresponds to (the cost function of) a sharpened version of the Riemannian metric given by the adjugate diffusion tensor [20, 19], which can be considered the preferred choice of metric in the Riemannian setting [39].

### 3.2. Path measures

Geodesic-based connectivity analysis combines a variety of curve shape measures with measures derived from the diffusion signal along the curve into a single path measure. This shape measure could be as simple as the Euclidean length, while more advanced shape measures such as local curvature and torsion are possible but used less frequently. The diffusion signal is encoded in the Finslerian length, representative of the total diffusivity, or the Finslerian speed, representative of the local diffusivity, or in a set of statistical measures such as the quantiles, mean, and standard deviation of the Finsler function evaluated along the geodesic. The heuristic definition of connectivity in terms of path measures is discussed in subsection 3.4 and in Appendix C.

The basic path measure used in the Finsler connectivity module is defined as

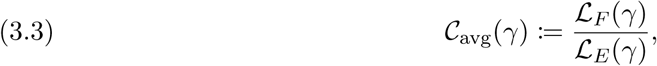

which is the Finslerian generalization of the most commonly used measure in Riemannian geodesic connectivity analysis [3, 28]. This measure can be interpreted as the average cost incurred along the geodesic, which is expected to be low for curves between two well-connected regions.

The second available path measure is defined as the largest local cost along a geodesic, given by

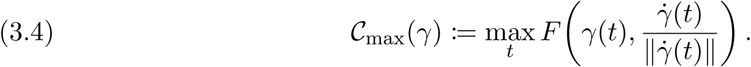

The 𝓒_max_ path measure was originally proposed in the Riemannian setting by Pechaud et al. [35]. Although this measure might be expected to be very sensitive to noise, it should be noted that it is based on the same (intrinsically smooth) geodesics as the 𝓒_avg_ measure, and is thus as stable as 𝓒_avg_. The 𝓒_max_ measure highlights geodesics that have continuously strong diffusivity along their paths, in contrast to the 𝓒_avg_ measure for which a locally weak diffusivity might be offset by very strong diffusivities further along the geodesic. Again, a low value of the path measure implies a high connectivity.

Details on the implementation of these measures can be found in Appendix A.

### 3.3. The Finsler connectivity module

The FCM typically returns two scalar maps—a distance map 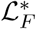 and the corresponding path measure map—based on an input diffusion-weighted data set that provides *F*, an optional mask image, and a label map containing labeled seed regions. The choice between *F*_old_ and *F*_new_ can be supplied as well, along with a choice of the path measures 𝓒_avg_ and 𝓒_max_ and further resolution and convergence parameters (e.g. the maximum order used in the spherical harmonic representation of *F*). The distance map provides at each position the shortest Finslerian distance to the seed region in accordance with (2.6), and the path measure map gives the path measure associated with the shortest geodesic connecting each point to the seed region. Geodesics can be computed if the internally generated tangent vector map is returned as well. The FCM is available at github.com/tomdelahaije/fcm, and can be used as a command-line tool.

### 3.4. Experimental design

We demonstrate the performance of the proposed Finsler-based connectivity analysis in two different settings: (1) a qualitative validation based on data from the WU-Minn Human Connectome Project (HCP), and (2) a quantitative network analysis of connectivity in autism spectrum disorder.

#### Qualitative validation

The qualitative Finsler connectivity validation experiments presented in subsection 4.1 are based on the pre-processed data of a single healthy subject in the WU-Minn Human Connectome Project (HCP) [45], released as part of the HCP 500 Subject Release. The data was acquired on a modified 3T Siemens scanner, with 1.25 mm isotropic voxels. The diffusion-weighted images are acquired on three shells (*b* = {1000,2000,3000} s/mm^2^) with 90 uniformly distributed gradient directions each, together with 18 baseline images. We express *F*_old_ and *F*_new_ in terms of spherical harmonics (maximum order 6), based on the *b* = 3000 s/mm^2^ shell. T1 data with an isotropic voxel size of 0.7 mm was also available. Further details on the acquisition protocol can be found on the HCP web site and in the references [45, 21, 2].

All maps based on the HCP data are seeded from manually selected voxels within the white matter, one voxel per bundle, based on the DTI white matter atlas by Catani and Thiebaut de Schotten [11]. Seeds are placed in four well-known major white matter bundles: the cingulum, the arcuate fasciculus, the corticospinal tract, and the splenium of the corpus callosum.

#### Network analysis in autism spectrum disorder

In order to provide an illustration of how the Finsler connectivity framework could be applied to population studies in a clinical setting, we present here a proof of concept network-based analysis of our Finsler connectivity approach, applied to autism spectrum disorder (ASD) data. To do so, we use a paradigm that is commonly used in network-based studies of brain connectivity. Specifically, we build a graph model of the brain, where graph nodes represent gray matter regions as defined through a FreeSurfer parcellation based on the Desikan–Killiany atlas [15]. The edge weights in this model represent the Finsler connectivity between pairs of FreeSurfer-defined gray matter regions. Once this network model is constructed for each subject, we compute its local efficiency measure. The local efficiency measure [37] is one of the many standard graph-theoretical measures that are commonly computed in network analysis studies, in order to quantitatively summarize the network structure. It is possible to compute other measures as well, but since this experiment is meant only as an illustration, we focus on a single measure (the ‘local network efficiency’) that has been previously implicated in ASD [38, 29]. Details on the connectivity analysis pipeline can be found in Appendix C.

Diffusion and structural MRI data were acquired from 69 typically developing male controls (TDC, age range: 8.0–14.4 years, mean: 10.7, standard deviation: 1.8) and 46 age-matched male autism spectrum disorder patients (ASD, age range: 8.1–14.1 years, mean: 10.7, standard deviation: 2.0). A t-test for difference in age between the two groups resulted in a p-value of 0.98. All imaging was performed using a Siemens 3T Verio scanner with a 32 channel head coil. Structural images were acquired on all subjects using an MP-RAGE imaging sequence (TR = 19 s, TE = 2.54 ms, TI = 0.9s, 0.8mm in-plane resolution, 0.9 mm slice thickness). Additionally, a HARDI acquisition was performed using a monopolar Stejskal-Tanner diffusion-weighted spin-echo, echo-planar imaging sequence with the following parameters: TR = 14.8 s, TE = 110 ms, 2 mm isotropic resolution, *b* = 3000 s/mm^2^, with 64 gradient directions and with two baseline images. The diffusion-weighted images of each subject were filtered using a joint linear minimum mean squared error filter to suppress Rician noise [43]. Eddy current correction was performed using registration of each volume to one of the baseline images. The same data was used in a study by Caruyer and Verma [10].

## 4. Results

### 4.1. Qualitative validation

All connectivity and path measure maps are shown in radiological convention. Path measure maps are shown using a temperature color map, where dark blue indicates a low connectivity/high path measure, and bright red indicates high connectivity/low path measure. The used color maps are shown on the right of each figure. Note that the results for the different path measures 𝓒_max_ and 𝓒_avg_ cannot be compared quantitatively, and that connectivity is strongest in regions that are connected to a seed point, typically leading to comparable high connectivity values within a *superset* of the bundle in which the seed is placed.

#### Cingulum

The first major bundle we consider is the cingulum, which consists of a set of fibers that project from the cingulate gyrus to the entorhinal cortex. In the work of Melonakos et al. [31] geodesic tractography was successfully used to trace this bundle, so we can expect our geodesic-based connectivity analysis to produce reasonable results for seeds placed in this bundle. The distance map from which geodesics can be computed is shown in Figure 2 for both *F* = *F*_old_ and the newly proposed *F*_new_-based metric. The Finslerian distance increases as expected from the seed outward, and the boundaries between white matter, gray matter, and cerebral spinal fluid can be identified very roughly. It is however difficult to judge the relative merit of the different metrics from these maps alone. Because all the connectivity measures under investigation in this section are derived from the same 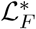 distance maps, and because the distance maps themselves provide little information, we omit the distance maps henceforth.

**Figure 2.**
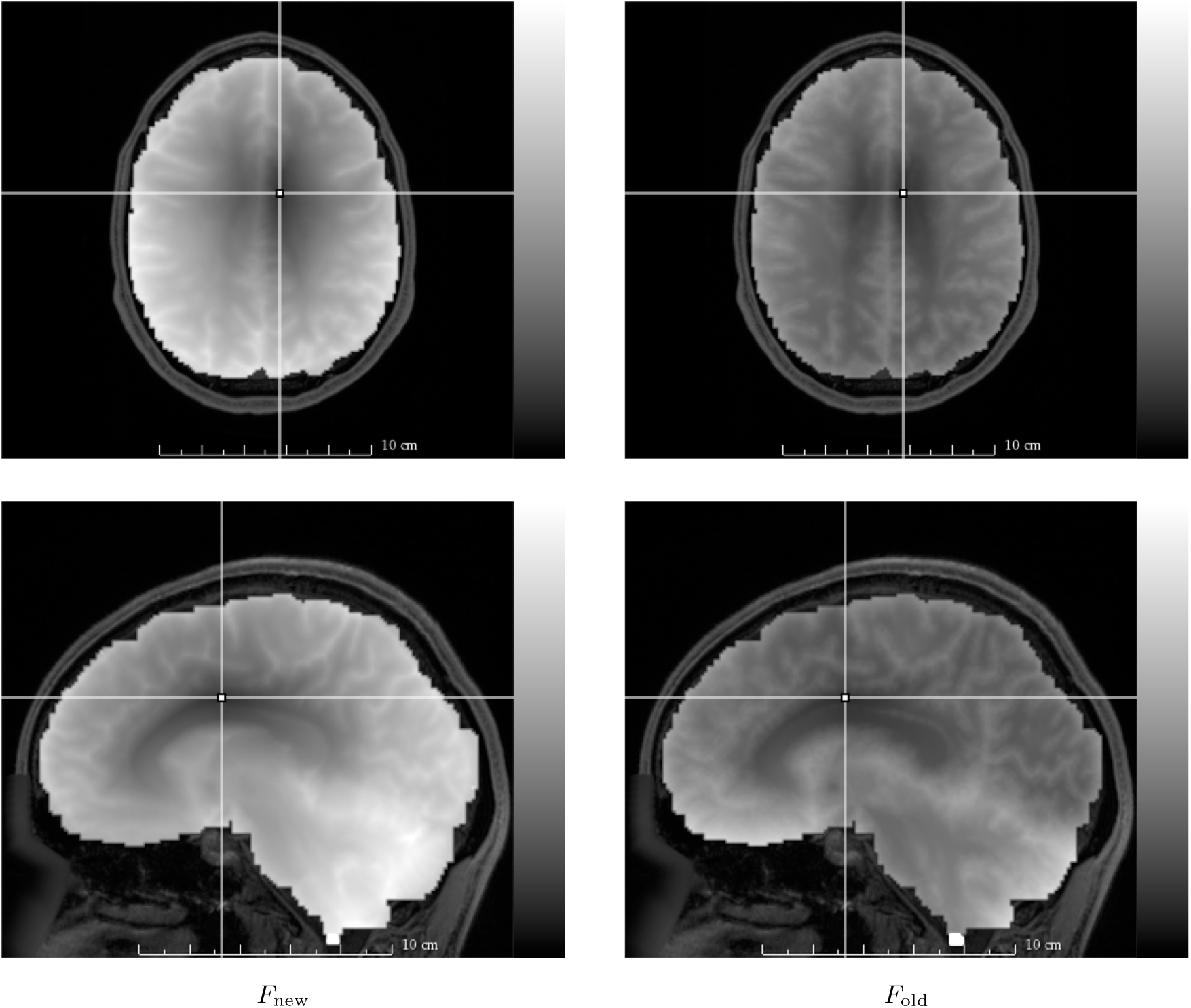
Axial (top row) and sagittal (bottom row) slices of distance maps 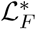 (2.6), for F = F_new_ (left column, (3.2)) and F = F_old_ (right column, (3.1)) seeded at a single voxel in the cingulum (annotated, point) of the HCP data set. Greater brightness encodes a greater distance. Other than minor differences in contrast, there are no conspicuous differences between the two choices for F deducible from these maps.

The connectivity maps shown in Figures 3 and 4 provide more information. First, a high-level comparison between Figures 3 and 4 reveals that the 𝓒_max_ connectivity measure (3.4) provides much more anatomical detail than the 𝓒_avg_ connectivity measure (Eq. (3.3)). In Figure 3 we see furthermore that the standard 𝓒_avg_ path measure map seeded in the cingulum bundle leaks into the corpus callosum, and to large sections of the posterior part of the brain. This effect is much less in the 𝓒_max_ map shown in Figure 4. Overall, the 𝓒_max_ map is much more specific to the known anatomy of the cingulum bundle, which allows us to perform a more detailed comparison between the *F* = *F*_old_ metric and our newly proposed *F* = *F*_new_ metric. We see that the former does result in leakage from the cingulum into the posterior parts of the corpus callosum (the splenium), and from there into large sections of posterior white matter. The choice *F* = *F*_new_ results in a path measure map that closely follows the known anatomy of the cingulum bundle, without leaking into the corpus callosum.

**Figure 3.**
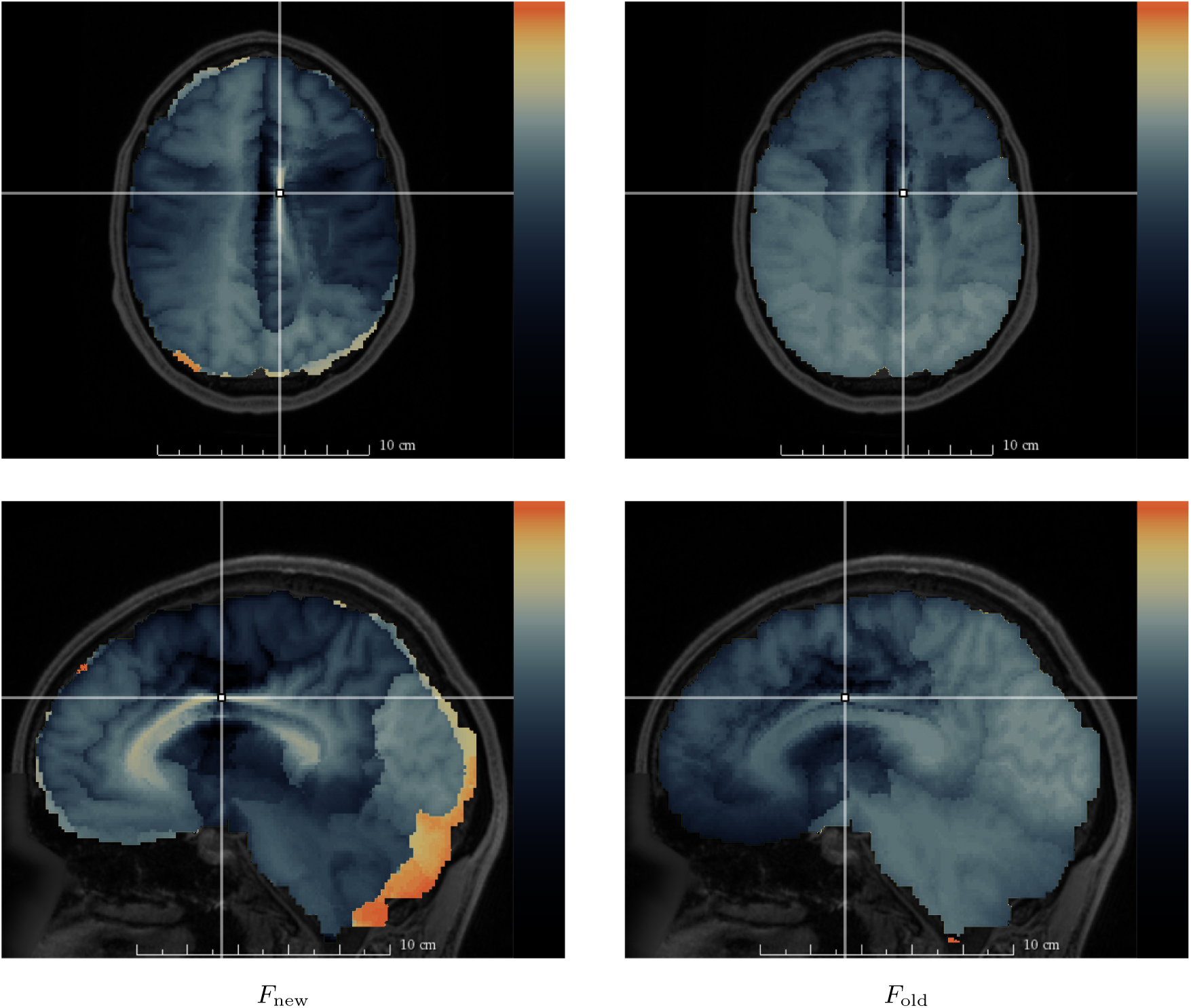
Axial (top row) and sagittal (bottom row) slices of maps based on the 𝓒_avg_ path measure (3.3), derived from the data shown in Figure 2 (seeded in the cingulum). The left column shows the results for the newly proposed F = F_new_ metric (3.2), and the right column shows results for the F = F_old_ metric (3.1). Bright red voxels are strongly connected to the seed region (white) according to the used path measures, while dark voxels are weakly connected. The displayed maps provide more anatomical detail than the distance maps in Figure 2, but appear to wrongly ascribe a high connectivity from the cingulum to the corpus callosum and to the posterior part of the brain. The choice F = F_new_ suffers from artificially high values at the edges, due to errors in the mask.

**Figure 4.**
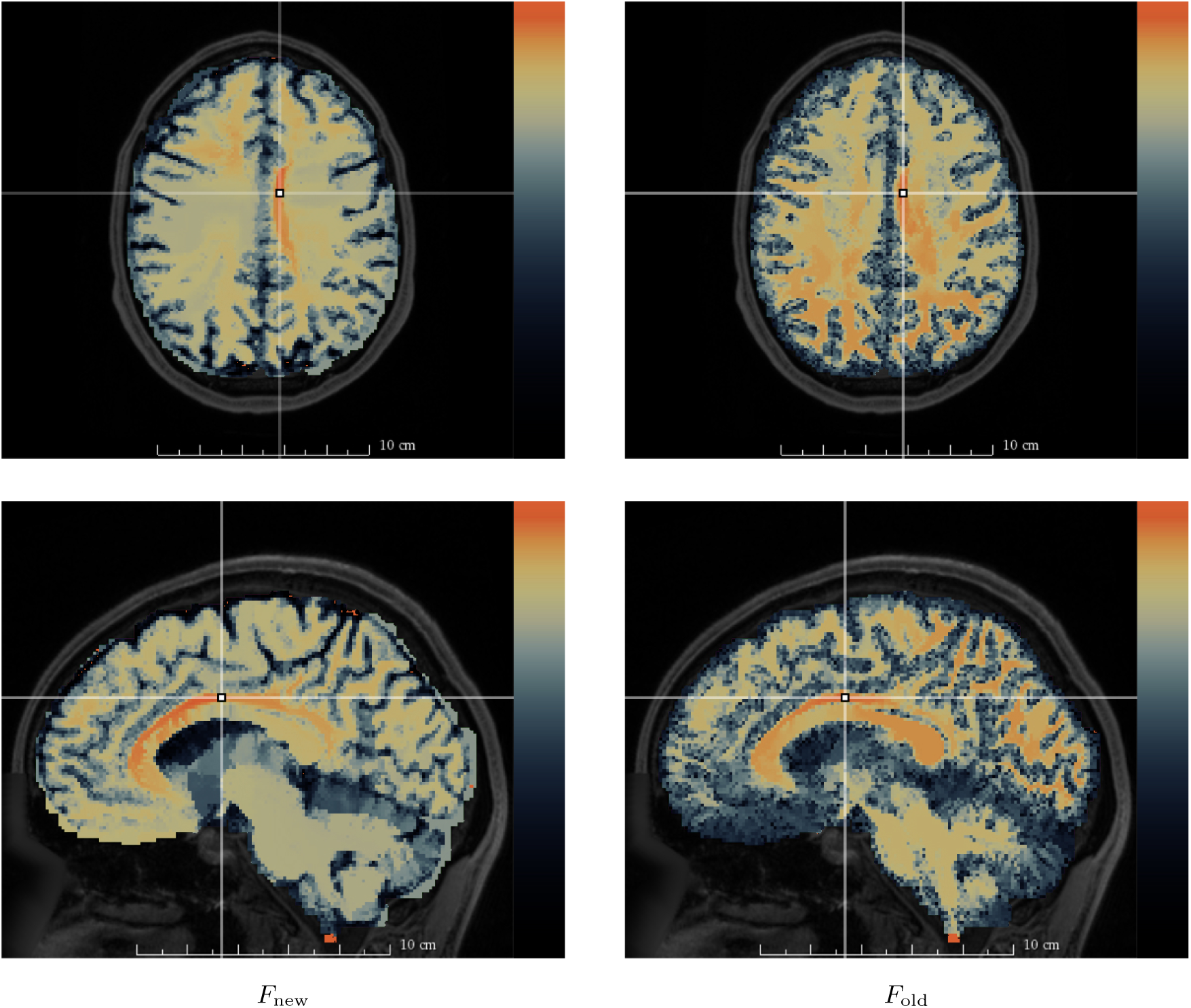
Axial (top row) and sagittal (bottom row) slices of maps based on the 𝓒_max_ path measure (3.4), derived from the data shown in Figure 2 (seeded in the cingulum). The left column shows the results for the newly proposed metric F = F_new_ (3.2), and the right column shows results for the F = F_old_ metric (3.1). We observe again that anatomical features are more distinguished compared to Figure 2, but also compared to Figure 3. Note that the F_old_ metric leads to much more significant ‘leakage’ into the corpus callosum. The 𝓒_max_ path measure shown in this figure is largely unaffected by masking errors, cf. Figure 3.

Finally we note that all maps suffer to a certain extent from errors in the mask, which can cause erroneous and problematic high path measure values near the boundaries of the mask. These false positives are clearly visible at the edges in Figure 3 (left) and near the brainstem in Figure 4 (bottom). With the 𝓒_avg_ measure these errors can propagate throughout the brain, which is particularly grievous in combination with the new *F*_new_-based metric. With the 𝓒_max_ path measure this propagation is however completely suppressed, which means the errors remain localized.

#### Arcuate fasciculus

Maps of the two path measures, 𝓒_avg_ and 𝓒_max_, are shown for the arcuate fasciculus in Figures 5 and 6. The arcuate fasciculus is a functionally important bundle, involved in aspects of language processing. It connects frontal cortical areas with the superior temporal gyrus. It also includes connections to the inferior parietal lobe. The maps for this bundle very clearly highlight a major issue with the widely used 𝓒_avg_ measure: despite the seed being placed in the right hemisphere, the left hemisphere shows an overall *stronger* connection to the seed than the voxels in the right hemisphere. This problem is to a large extent resolved with the introduction of the 𝓒_max_ measure. For the remaining bundles we show only results with the 𝓒_max_ measure, though it should be noted that the 𝓒_avg_ measure can produce good results *locally* as shown in the top-left map in Figure 5.

**Figure 5.**
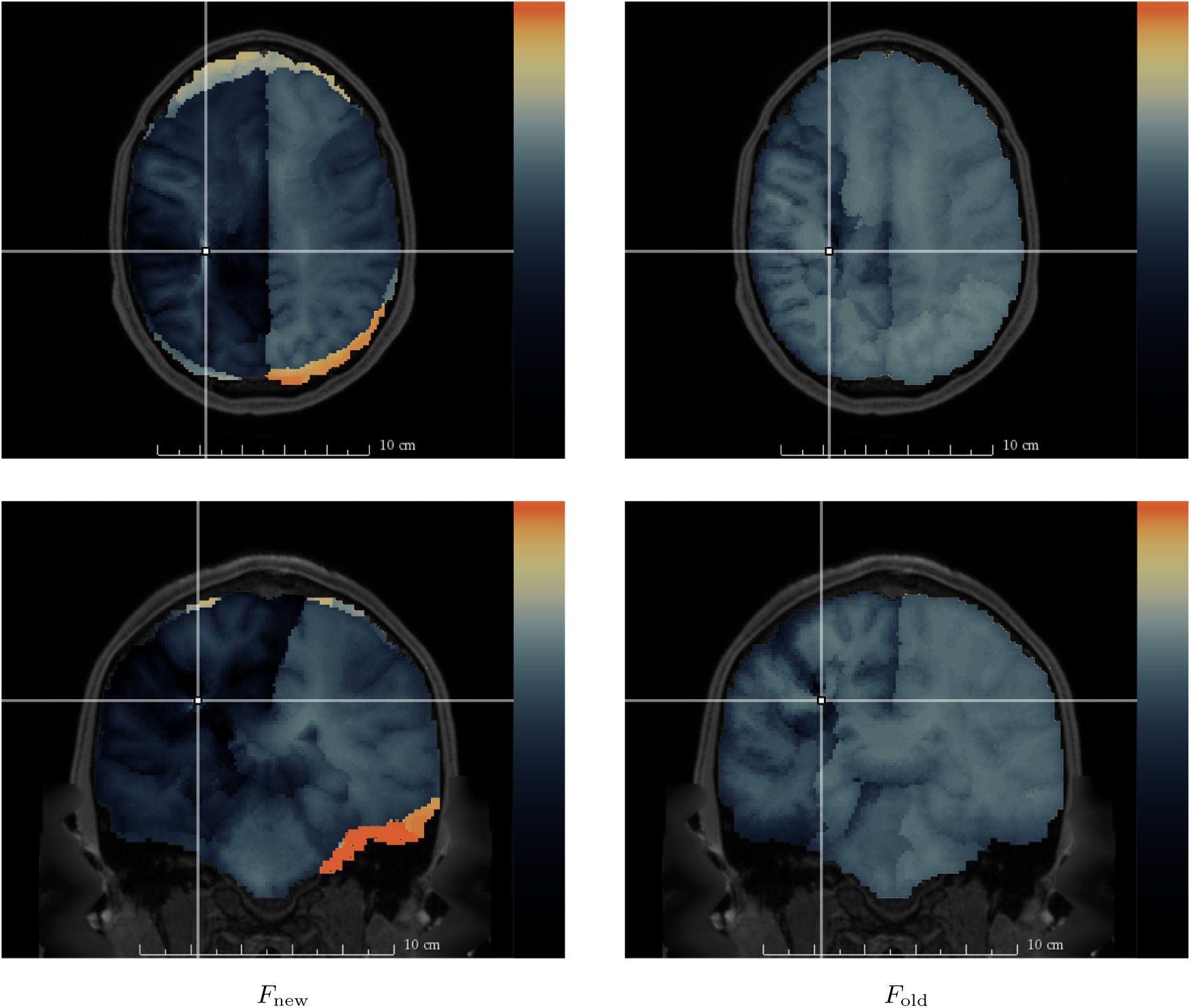
Axial (top row) and coronal (bottom row) slices of 𝓒_avg_-based maps (3.3) for F = F_new_ (left column, (3.2)) and F = F_old_ (right column, (3.1)) seeded in the arcuate fasciculus (annotated, point) of the HCP data set. Connections over the corpus callosum typically have large average diffusivities, which results in the undesired high values in the unseeded hemisphere visible for both metrics.

**Figure 6.**
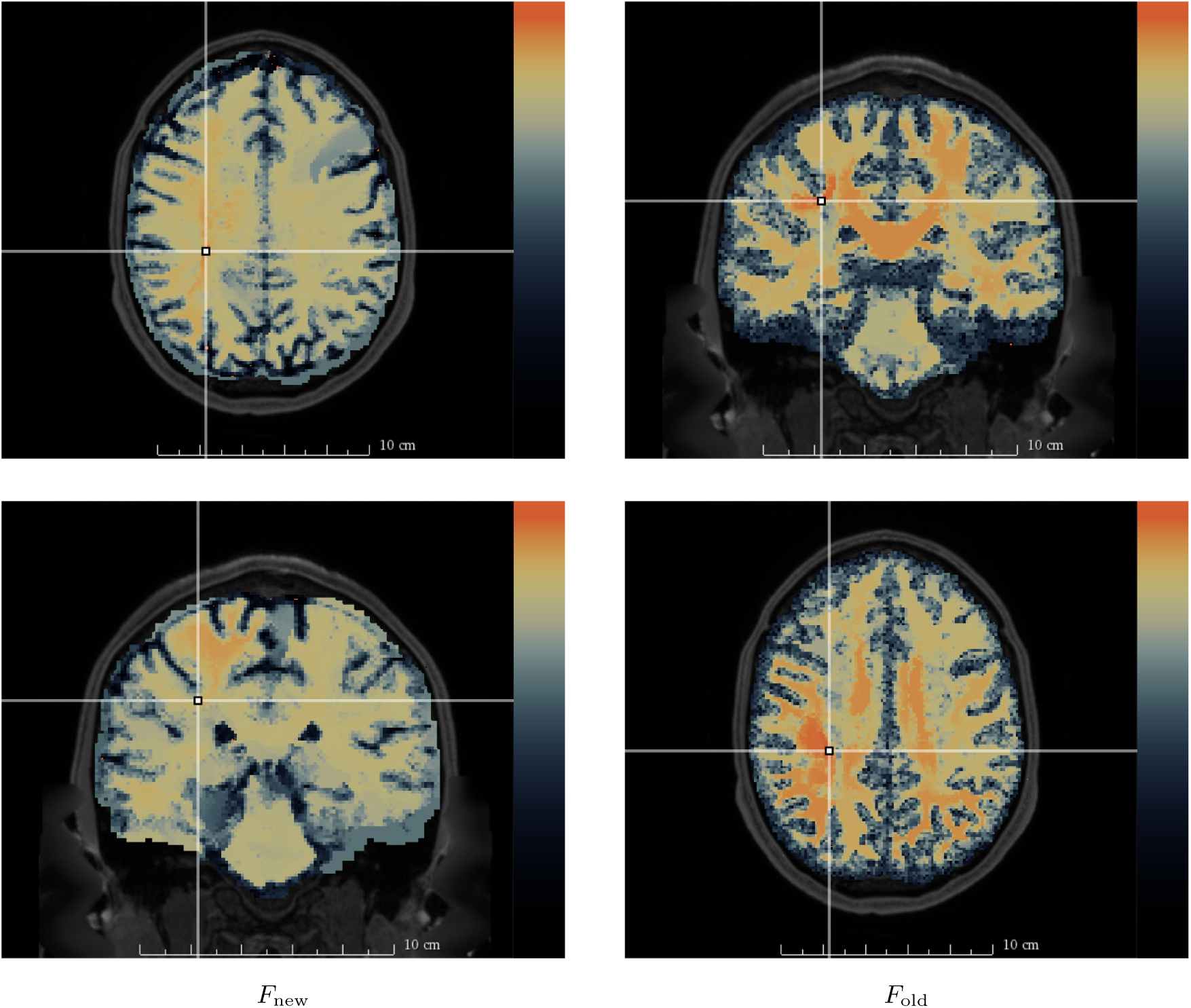
𝓒_max_-based maps (3.4) for F = F_new_ (left column, (3.2)) and F = F_old_ (right column, (3.1)) corresponding to Figure 5 (seeded in the arcuate fasciculus). Unlike the corresponding 𝓒_avg_-based maps shown in Figure 5, the 𝓒_max_ path measure does not have a bias for cross-hemispheric connections.

Again, a comparison between the 𝓒_max_-based maps with *F* = *F*_old_ and *F* = *F*_new_ reveals a superior performance of the new *F*_new_-based metric (barring the artifacts due to masking errors). With the *F*_old_-based metric, there is again leakage into the corpus callosum and then into the opposite hemisphere, which is not observed with the *F*_new_-based metric. As it is difficult to judge the reconstruction of the arcuate fasciculus from 2D slices, we show a 3D reconstruction of a thresholded path measure map obtained with *F* = *F*_new_ in Figure 7, which shows a fairly complete reconstruction of the regions connected to and including the arcuate fasciculus with minimal leakage into other bundles. This reconstruction was impossible with *F* = *F*_old_.

**Figure 7.**
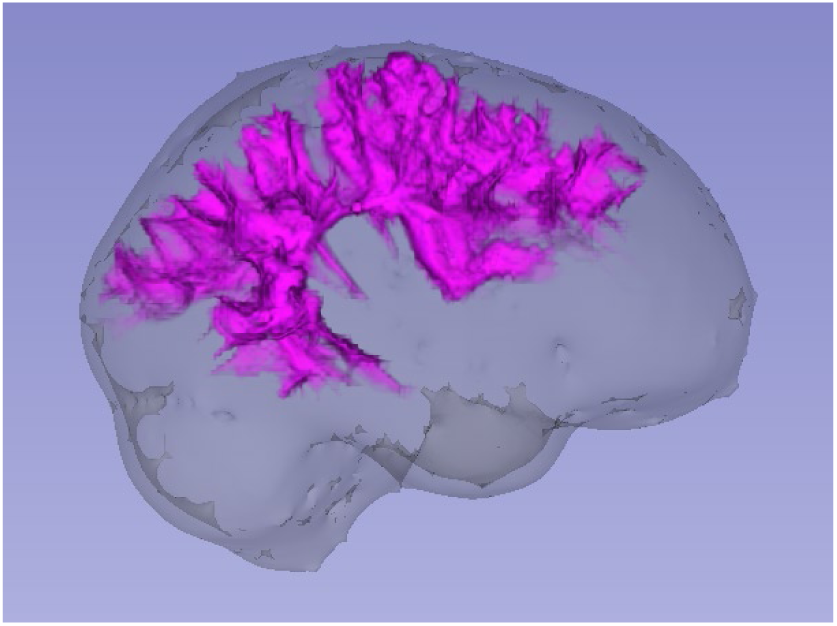
A left-right view of a rendering of the regions connected to the arcuate fasciculus obtained by thresholding the 𝓒_max_-based map (3.4) (with F = F_new_) shown in Figure 6. The rendered segmentation envelops the arcuate fasciculus, is entirely contained within the right hemisphere, and contains further offshoots of structures connected, to the seeded bundle. Attempts to reconstruct a similar segmentation using the 𝓒_max_-based map with F = F_old_ expectedly included, large regions contralateral to the seeded bundle, without capturing the full structure of the arcuate.

#### Corticospinal tract

The next tract we examine is the corticospinal tract (CST), which is a major fiber tract that conducts sensorimotor signals between the cortex and the spinal cord. 𝓒_max_ path measure maps obtained with a single voxel seed in the CST are provided in Figure 8. These maps show some of the advantages and disadvantages of the different metrics. The traditionally used *F*_old_ metric appears to be better equipped to avoid leakage into the opposite hemisphere, at the level of the brain stem, and specifically at the pons, where pontine crossing fibers may cause the connectivity to cross the midline into the other hemisphere [46]. To see this, compare the images in the bottom row of Figure 8. This result is anatomically incorrect and is a well-known issue with many tractography algorithms [46]. While the *F*_new_ metric is more sensitive to this issue, the regions of high connectivity extend much further in the superior direction towards the cortex, better reproducing the known fiber fanning in this region. Furthermore, *F*_new_ based connectivity of the CST shows only a minor leak into the corpus callosum, while the *F*_old_ metric produces a much more extensive leak into the posterior part of the corpus callosum and into the opposite hemisphere.

**Figure 8.**
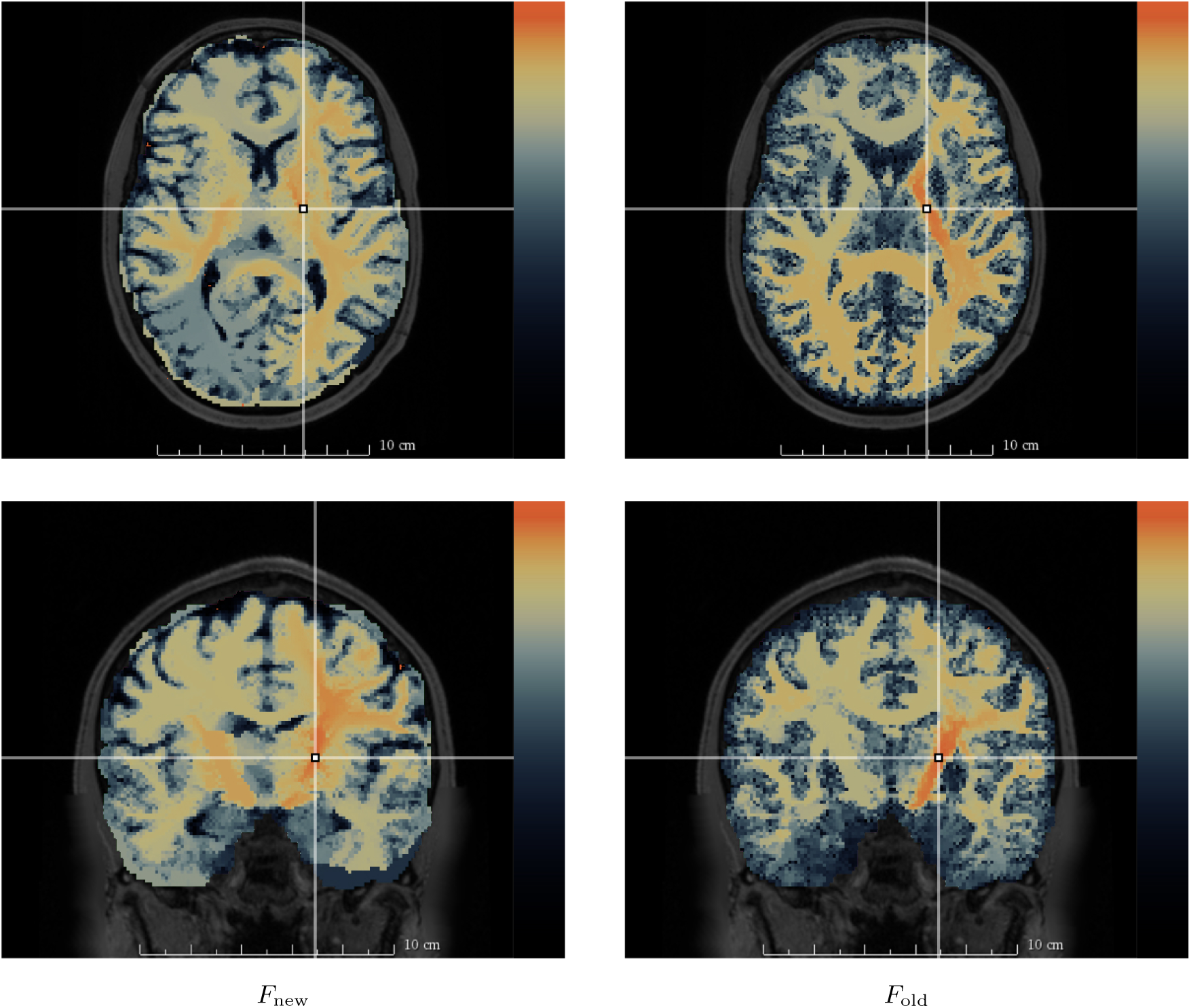
Axial (top row) and coronal (bottom row) slices of 𝓒_max_-based maps (3.4) for F = F_new_ (left column, (3.2)) and F = F_old_ (right column, (3.1)) seeded in the corticospinal tract (annotated point) of the HCP data set. For F = F_new_ connectivity leaks to the opposite hemisphere at the level of the brain stem where pontine fibers cross into the other hemisphere, while the regions of high connectivity fan and extent superiorly to a greater degree than for F = F_old_.

#### Corpus callosum

The last bundle we consider is the corpus callosum, a massive white matter highway that connects the two hemispheres. The 𝓒_max_-based path measure maps for the *F*_old_ and *F*_new_ metrics were seeded in the splenium (posterior part) of the corpus callosum, as shown in Figure 9. With the *F*_new_ metric, we observe that the path measure map remains concentrated in the posterior parts of the corpus callosum, while with the *F*_old_ metric it progresses much further in the anterior direction, which should not be happening with a seed located posteriorly.

**Figure 9.**
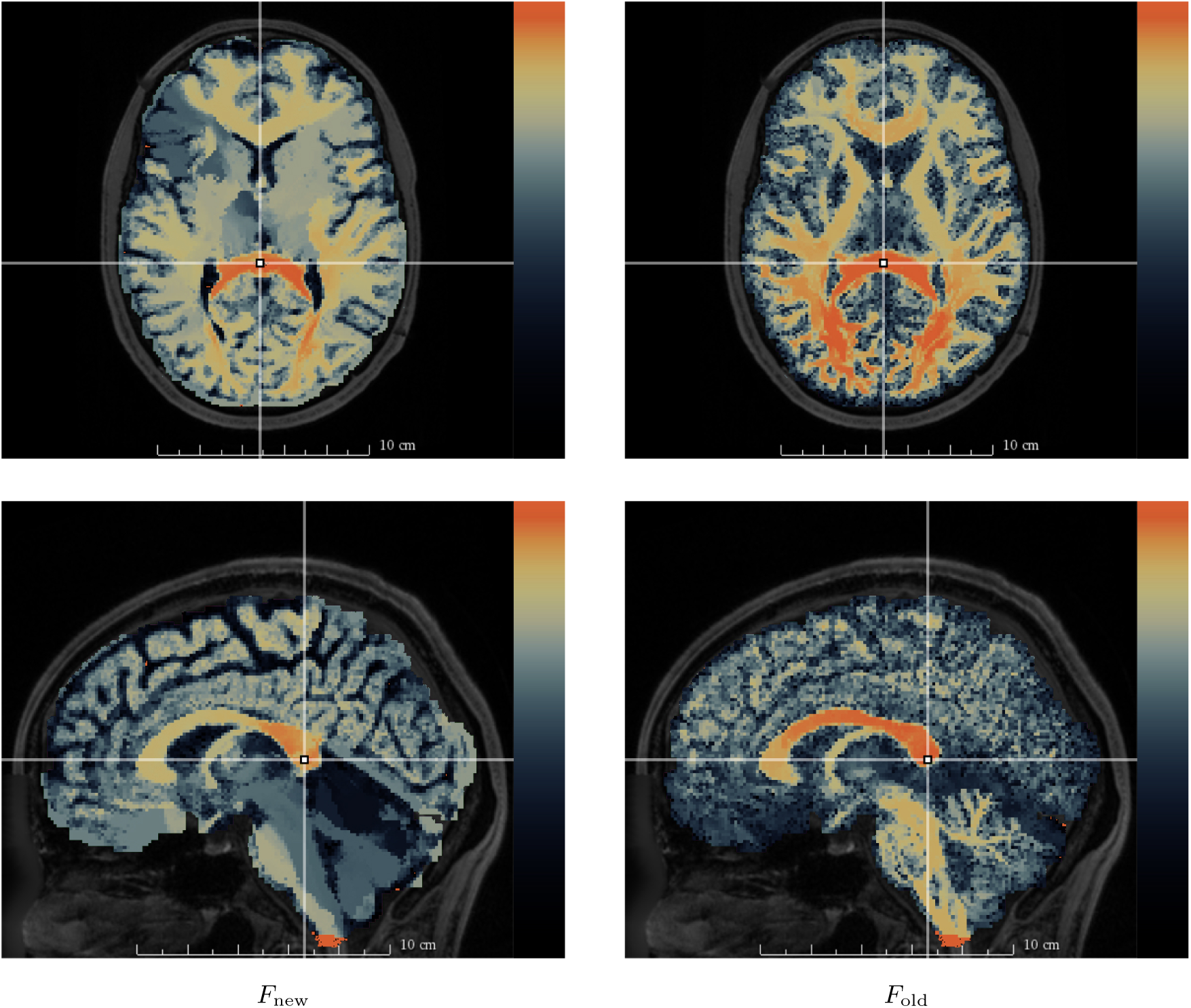
Axial (top row) and sagittal (bottom row) slices of 𝓒_max_-based maps (3.4) for F = F_new_ (left column, (3.2)) and F = F_old_ (right column, (3.1)) seeded in the splenium of the corpus callosum (annotated, point) of the HCP data set. The F_new_-based metric follows anatomy more closely than the F_old_-based one, exemplified here by the high values of the latter observed in the frontal part of the corpus callosum.

### 4.2. Network analysis in autism spectrum disorder

We performed the network analysis study with the newly proposed *F*_new_-based Finsler metric, using three different maximum orders for the spherical harmonic representation of the ODF. For comparison we also repeated the experiment, for the same three different spherical harmonic orders, using the *F*_old_-based metric proposed by Melonakos et al. [32] (3.1). The resulting p-values are presented in Table 1. All experiments used the 𝓒_max_-based connectivity measure.

**Table 1.** p-values for the MANOVA comparison between the local efficiency vectors of the control and ASD groups. Column heading F_new_ indicates connectivity computed using F = F_new_ metric, whereas F_old_ denotes the metric proposed by Melonakos et al. [32]. Connectivity matrices were computed using a spherical harmonic representation of the diffusion ODF with, three different orders: 2, 4 and 6. All connectivity measures were computed with, the 𝓒_max_ path, measure. Since a total of six tests were performed, the Bonferroni-corrected threshold for significance was 0.05/6 ≈ 0.0083. p-values below this threshold (shown in **bold**) indicate rejection of the null hypothesis that there is no difference between the two groups.

| Order | $F_{\text{new}}$ | $F_{\text{old}}$ |
| --- | --- | --- |
| 2 | **0.0035** | **0.0061** |
| 4 | **$1.7 \times 10^{-4}$** | 0.0087 |
| 6 | 0.035 | 0.030 |

## 5. Discussion

In this work we described the basic ideas of Finsler geometry for the purpose of geodesic connectivity analyses, and introduced the open-source Finsler connectivity module (FCM) for Finsler geodesic tractography and connectivity studies. The FCM is based on the `finslertract` project of Antonio Tristán-Vega (nitrc.org/projects/finslertract), and includes changes that reflect recent advances in Riemannian geodesic tractography [19, 35].

The connectivity analysis capacities of the module were primarily evaluated on Human Connectome Project (HCP) data, with results depending heavily on the choice of Finsler function and path/connectivity measure.

We have considered two different Finsler metrics, one based on a newly proposed Finsler function *F*_new_ derived from work by Fuster et al. [19] in the Riemannian setting, and the *F*_old_-based metric originally proposed by Melonakos et al. [32], which is the one typically used in literature. While a visual assessment of the distance maps obtained with each approach is not informative (see e.g., Figure 2), the corresponding path measure maps highlight some interesting differences. In particular, the maps obtained using the Finsler function *F* = *F*_new_ are more faithful to the known anatomy of tracts, as illustrated by the examples in subsection 4.1. Although both the *F*_new_- and *F*_old_-based maps suffer some ‘leakage’ problems, i.e., high connectivity values spreading to nearby but unrelated tracts, the *F*_new_-based metric is much more robust to this issue compared to the *F*_old_-based metric. This is especially clear near the corpus callosum, as can be seen in the cingulum results shown in Figure 4. Thus, in addition to its more rigorous theoretical foundation, the *F*_new_ Finsler function typically results in connectivity maps which are anatomically more reliable.

In contrast to the *F*_old_-based Finsler metric, the new metric is designed to correspond to a theoretically well-founded Riemannian metric [19], cf. Appendix B. However, the fact that the generalized Finslerian metrics used here are still defined in an ad hoc manner remains a significant issue with the interpretation of Finsler geometrical analyses. Given the improved results obtained with the relatively simple *F*_new_ Finsler function, it will be worthwhile to investigate the application of the fundamental results presented in the work of Dela Haije et al. [14] to the current analysis pipeline in future work.

In subsection 3.2 and Appendix A we explained how the 𝓒_max_ measure used in the Riemannian setting by e.g. Pechaud et al. [35] can be applied in the Finslerian setting. This measure is based on the ‘weakest link’ of the geodesic. That is, geodesics between strongly connected points should have a continuously low cost along the entire tract, while even small regions of high cost along the geodesic are taken to significantly decrease the likelihood of the points being structurally connected. Compared to the more common 𝓒_avg_ measure, which considers the average of some diffusivity measure along a geodesic, the 𝓒_max_ path measure has a number of important advantages, both theoretical and practical.

Primarily, the 𝓒_max_ measure associates a relatively low connectivity to geodesics taking shortcuts, a notorious problem of geodesic tractography [41, 23]. In the same vein, provided a sufficiently fine spatial resolution, low connectivity is associated with geodesics that jump from one fiber bundle to another across a small region with high cost. Because of this, in practice, regions of high connectivity tend to concentrate much more on the seeded fiber bundles, which again significantly reduces leakage artifacts. One can appreciate this especially in the difference between the 𝓒_avg_ and 𝓒_max_ path measure maps seeded in the arcuate fasciculus (Figures 5 and 6), where the consistently low cost in the corpus callosum results in an above average 𝓒_avg_-based connectivity for all geodesics that cross the corpus callosum to the other hemisphere. This in fact highlights another theoretical advantage; the 𝓒_max_ measure has the very natural property of being monotonic, i.e., it cannot increase in connectivity with distance along a path. Combined, these properties result in 𝓒_max_-based connectivity maps that are generally much closer to anatomy than maps produced using the 𝓒_avg_ measure.

However, the 𝓒_avg_ measure can lead to greater contrast than 𝓒_max_ at a *local* level, as can be seen in the left columns (*F*_new_-based metric) of the arcuate fasciculus results, Figures 5 and 6. 𝓒_max_-based maps typically have low homogeneous connectivity throughout the white matter, which essentially results in a default situation in which everything is (at least weakly) connected to everything else, while 𝓒_avg_-based maps can efficiently extract the local, more direct structural connections.

The large differences between the two path measures highlight an issue with the currently employed definitions. The different types of information captured by these measures make it clear that other path measures, or more likely combinations of various measures, can and should be developed to obtain a more complete characterization of the structural connectivity captured by the geodesics. Because the validation of connectivity measures is very challenging, a possible next step could be a structured inclusion of a complete set of descriptive measures, e.g. shape measures of increasing complexity, subject to natural constraints like scale and orientation invariance. The introduction of anatomical priors, which has become more common in recent works [42], could also be used to further improve geodesic-based connectivity analysis.

Finally, we have studied group differences in a graph-theoretical analysis of brain networks in autism spectrum disorder (ASD), where we found significant differences in the local network efficiency between the ASD group and the normal developing controls. These results depend on the spherical harmonics used to represent the Finsler function, Table 1, which needs to be chosen in a manner that balances angular resolution and sensitivity to noise. Differences in local network efficiency have been widely reported in previous works, and our results corroborate previous findings of abnormalities [38, 29]. Note that this analysis was not intended to be exhaustive, but rather to illustrate the application of our new connectivity framework in a standard brain network analysis setting. In future work, we will conduct a more thorough analysis of brain networks based on Finsler connectivity.

The main problem addressed only superficially in this work is validation of the connectivity measures. As with available alternatives, the absence of ground truth data makes it very challenging to validate and compare connectivity measures. In the near future this may become feasible, with the way being paved by recent efforts on validation of microstructural models through e.g. histology [40], and on validation of tractography by evaluating methods on simulated data with a known ground truth [33, 30]. Although outside of the scope of this work, a direct comparison of different connectivity measures in specific applications (such as population studies) could provide additional insight into the performance of the proposed measure.

In conclusion, we have presented a new tool, the Finsler connectivity module, that implements a connectivity pipeline for diffusion MRI. To our knowledge this is the first non-Gaussian geodesic connectivity method made available to the neuroimaging community in the form of a ready-to-use tool. Our implementation improves the existing framework used by e.g. de Boer et al. [13], as demonstrated in the presented HCP experiments. These improvements derive from a novel Finsler function defined in terms of the solid angle ODF, and from the consideration of more appropriate connectivity measures. In future work we will evaluate our approach with respect to the currently employed tractography-based connectivity methods.

## Appendix A Implementation details

In the pseudo-code below (Algorithm 1) we present a modification of Melonakos’ fast-sweeping algorithm [32], with the additional steps needed to compute the proposed path measures ((3.3) and (3.4)). In analogy with the notation for the distance map 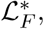 which gives the shortest distance between each point and the seed region Ω, we define 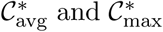 as the path measure maps that give for each point the path measure associated to the geodesic from that point to the considered seed region. Similarly, we define 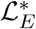 as the shortest Euclidean distance between each point and the seed region, and ***y***^∗^ as the tangent to the geodesic at each point. The algorithm is based on the observation that for a solution of (2.8), the Finslerian distance between ***x*** and ***x*** + ***y*** for ‖***y***‖ small is simply given by *F*(***x***, ***y***). The exact definition of the addition ***x*** + ***y*** requires the exponential map on the Finsler manifold, which goes beyond the scope of this paper—see e.g. the work by Bao et al. [5] for more information. In the FCM interpolation is done linearly [32], but alternative methods are possible.

## Appendix B. Derivation of *F*_new_

In the usual Riemannian framework for DTI [34, 27] the metric tensor ***g*** is defined by the inverse diffusion tensor, i.e., ***g*** = ***D***^−1^. A minor modification of this relation has recently been proposed on theoretical grounds [20, 19], in which a new metric ***g̃*** is defined by the adjugate diffusion tensor:

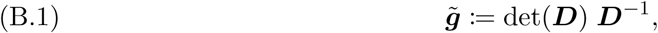

where det ***D*** is the determinant of the diffusion tensor ***D***. This relation between *F*_new_ and the adjugate *g̃* can be recognized in the exact expression for the normalized ODF [26]

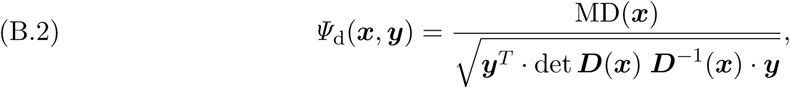

where MD is the mean diffusivity and ***y*** ∈ S^2^ is a direction unit vector. We note that the denominator in this equation is the norm of direction vector ***y*** in the Riemannian space equipped with the metric ***g̃***:

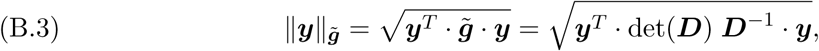

and combining (B.2) and (B.3) we find

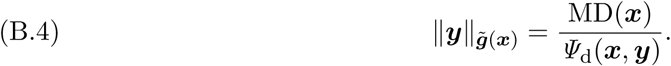

### Algorithm 1

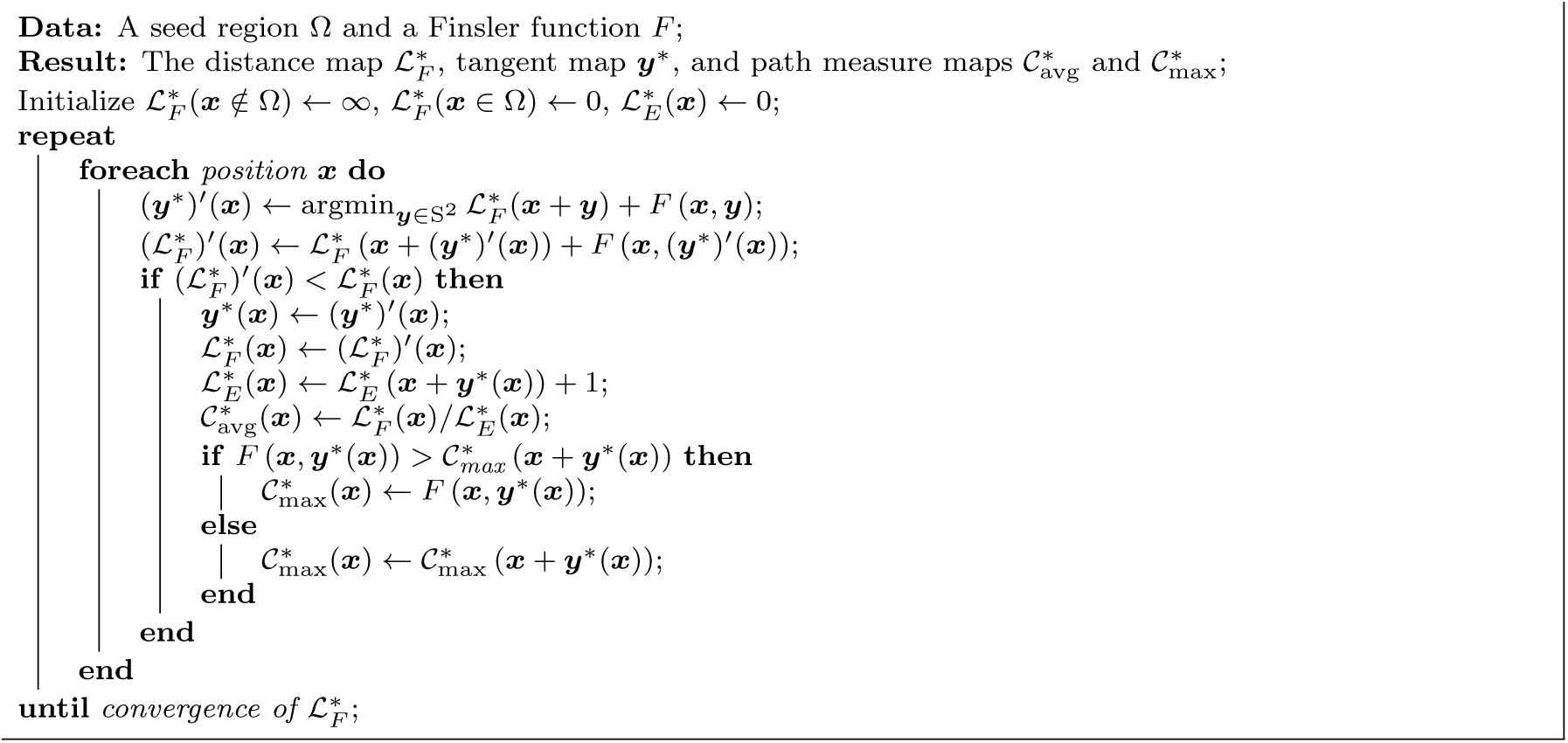
The fast-sweeping algorithm used to compute the maps 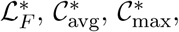 and ***y***^∗^, adapted from Melonakos et al. [32]. The implementation is based on the `finslertract` project of Antonio Tristán-Vega.

Replacing *Ψ*_d_ with the sharper *Ψ*_sa_ then produces the *F*_new_ metric proposed in (3.2).

## Appendix C. Connectivity analysis pipeline

In recent years, the view that the functional and structural systems of the brain can be modeled as complex networks has motivated a large amount of research on the application of graph theoretical concepts to brain network analysis [37, 9]. The standard graph network model of the brain consists of a set of nodes, which represent a partitioning of the cortex and other gray matter structures. These nodes are connected via a set of edges, or links, that represent structural and/or functional connections between gray matter partition units.

Such a graph model of the brain’s network organization can be constructed from a variety of imaging modalities such as structural MRI, diffusion MRI, functional MRI, or EEG/MEG. In this setting, a characterization of the organization of the different computational nodes and the functional interaction between them is achieved via graph theoretical analysis [37, 9]. This appendix describes the details of the pipeline used to perform the group analyses reported in this work, covering the cortical surface parcellation, the connectivity measure computation, and the final statistical analysis.

### FreeSurfer parcellation

FreeSurfer is a freely available software toolbox that reconstructs mesh-based models of the cortical surface [12, 17], and provides a parcellation of the cortex into neuro-anatomical areas using both the geometrical model of the cortical surface and neuro-anatomical convention [16].

For the present set of experiments, we computed a FreeSurfer parcellation for each subject based on the Desikan–Killiany atlas [15], which resulted in a total of 86 cortical and subcortical gray matter regions. FreeSurfer parcellations are initially defined in each subject’s T1 standard space. In order to allow for these gray matter regions of interest to be used as seed regions for Finsler connectivity analysis, we registered the FreeSurfer parcellation to the diffusion MRI space using non-linear registration.

### Connectivity matrix construction

The path measures discussed in subsection 3.2 are length-based, i.e., low values indicate a short geodesic distance, which implies a high connection strength. From these path measures we can heuristically derive a new measure that signifies a general notion of connectivity as follows.

For each subject in our study, we constructed an 86×86 connectivity matrix ***C***, whose *ij*-th entry *C_ij_* represents the connectivity between the *i*-th and *j*-th FreeSurfer regions. Each entry *C_ij_* is computed as follows. First, seeding in the *i*-th region, we run our Finsler connectivity algorithm to produce a path measure map for the entire brain. Then, the values for the path measure in the spatial extent of the *j*-th region are averaged for each *j* ≠ *i*, which results in the average path measure values *m*_*i*→*j*_. This process is repeated for all seed regions *i*. Finally, we construct the connectivity matrix ***C***:

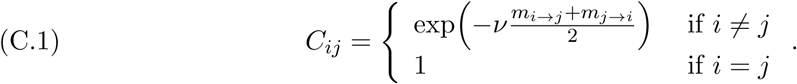

Note that ***C*** is a symmetric matrix by construction, so each pair of regions has a uniquely defined connectivity value. The scaling parameter *ν* is set to 0.1 for the presented experiments, producing a reasonable spread of connectivity values over the range [0,1] with small values indicating a low connection strength.

As an example, Figure 10 shows a coronal slice through a path measure map computed on one of the TDC subjects. This map was seeded in the left caudal middle frontal gyrus region, as defined by a FreeSurfer segmentation using the Desikan–Killiany atlas as described above. The seed voxels that intersect this particular coronal slice are shown in white with a black outline. The path measure map itself is shown in a modified temperature color map, such that bright red colors indicate a low path measure and a dark blue color indicates a high path measure value. Thus, in order to compute *m*_*i*→*j*_, where *i* indicates the caudal middle frontal region, and *j* indicates any of the other FreeSurfer-defined regions, we average the values of this path measure map over the voxels that comprise FreeSurfer region *j*. Intuitively, regions that are well-connected to region *i* will result in low values for *m*_*i*→*j*_, which in turn will lead to higher connectivity values *C_ij_* according to (C.1).

**Figure 10.**
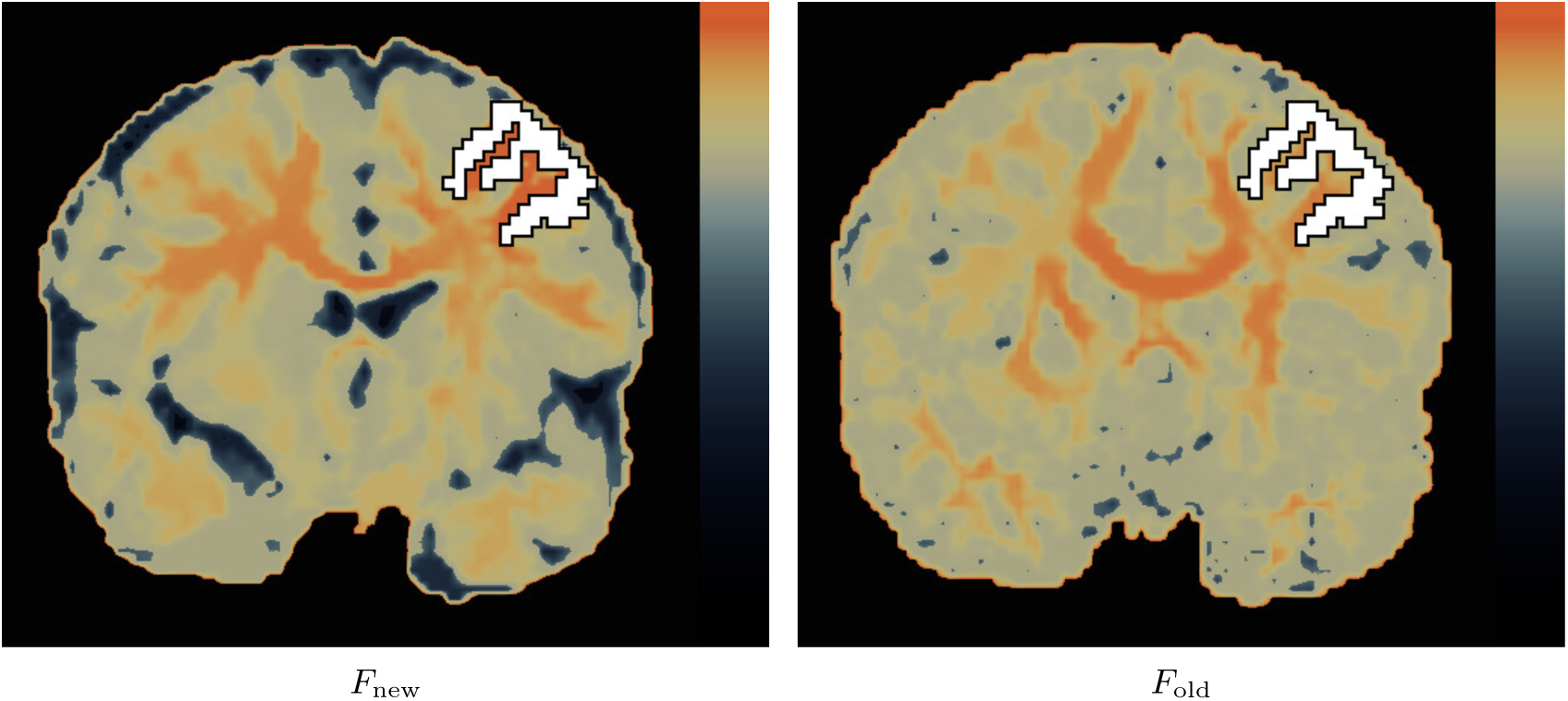
A comparison of 𝓒_max_ path measure maps, using the F_new_- and F_old_-based metrics on data from one of the TDC subjects. These maps are seeded in the left caudal middle frontal gyrus region, as defined by FreeSurfer. The seed voxels that intersect this particular coronal slice are shown in white. Bright red voxels are strongly connected to the seed region according to the used path measures, while dark voxels are weakly connected.

In addition to illustrating the concept of a path measure map and the corresponding computation of *m*_*i*→*j*_, Figure 10 also provides an initial comparison between the *F*_new_- and the *F*_old_-based metrics. With the caudal middle frontal gyrus as a seed region, the newly proposed *F*_new_-based metric results in the recovery of the known transcallosal connectivity to the opposite hemisphere (Figure 10, left). In contrast, the *F*_old_-based metric produces a map (Figure 10, right) that has drawbacks similar to known artifacts observed with tractography on DTI data. In particular, the connectivity does not reach the contralateral cortical regions, as it appears to stop in the well-known region of three-way crossings between the corpus callosum, the corticospinal tract and the superior longitudinal fasciculus. On the other hand, connectivity appears to ‘leak’ into the corticospinal tract of both hemispheres, as well as into other white matter tracts that do not have direct anatomical connectivity with the seed region. Furthermore, the *F*_new_-based metric correctly identifies CSF areas, such as the ventricles, and assigns these a high path measure (low connectivity). In contrast, the *F*_old_-based metric does not detect CSF and as a result propagates connectivity through CSF regions, which is clearly anatomically incorrect. The differences between the two metrics is addressed in more detail in subsection 4.1.

### Computation of local efficiency and statistical analysis

Once the connectivity matrix ***C*** is computed for each subject, we compute its local efficiency measures using the Brain Connectivity Toolbox [37]. The local efficiency of a network is a quantity computed at each node of the network, such that it quantifies the network’s resistance to failure at the local scale. In other words, it quantifies the importance of a graph node by measuring how well information is exchanged by the immediate neighbors of the node when it is removed. Thus, in the present experiment, this computation results in a 1 × 86 vector, such that its *i*-th element corresponds to the local efficiency measure of the *i*-th FreeSurfer region.

In the present experiment, we are interested in performing a statistical test for a group difference between the TDC and ASD groups of subjects based on their local efficiency vectors. To reduce the number of multiple comparisons, we do not test each region individually but perform a one-way multivariate analysis of variance (MANOVA) to compare the mean vectors for the two groups. This is a statistical test for the null hypothesis that the mean local efficiency vectors of the TDC and ASD groups are the same. If we can reject this null hypothesis, we conclude that the two groups differ in terms of their local efficiency measure, although the test does not identify specific regions that may be responsible for this difference.

## Acknowledgments

Data were provided in part by the Human Connectome Project, WU-Minn Consortium (Principal Investigators: David Van Essen and Kamil Ugurbil; 1U54MH091657) funded by the 16 NIH Institutes and Centers that support the NIH Blueprint for Neuroscience Research; and by the McDonnell Center for Systems Neuroscience at Washington University. Andrea Fuster and Tom Dela Haije would like to acknowledge their hosts at the Laboratory for Mathematics in Imaging at the Brigham and Women’s hospital, Harvard Medical School, for their hospitality.

